# The Dynamics of Attention Shifts Among Concurrent Speech in a Naturalistic Multi-Speaker Virtual Environment

**DOI:** 10.1101/626564

**Authors:** Keren Shavit Cohen, Elana Zion Golumbic

## Abstract

Focusing attention on one speaker on the background of other irrelevant speech can be a challenging feat. A longstanding question in attention research is whether and how frequently individuals shift their attention towards task-irrelevant speech, arguably leading to occasional detection of words in a so-called unattended message. However, this has been difficult to gauge empirically, particularly when participants attend to continuous natural speech, due to the lack of appropriate metrics for detecting shifts in internal attention. Here we introduce a new experimental platform for studying the dynamic deployment of attention among concurrent speakers, utilizing a unique combination of Virtual Reality and Eye-Tracking technology. We created a Virtual Café in which participants sit across from and attend to the narrative of a target speaker. We manipulate the number and location of distractor speakers, manifest as additional patrons throughout the Virtual Café. By monitoring participant’ eye-gaze dynamics, we studied the patterns of overt shifts of attention among the concurrent speakers as well as the consequences of these shifts on speech comprehension.

Our results reveal important individual differences in the gaze-pattern displayed during selective attention to speech. While some participants stayed fixated on a target speaker throughout the entire experiment, approximately 30% of participants frequently shifted their gaze toward distractor speakers or other locations in the environment, regardless of the severity of audiovisual distraction. Critically, the tendency for frequent gaze-shifts negatively impacted comprehension of the target speaker. We also found that gaze-shifts occurred primarily during gaps in the acoustic input, suggesting they are prompted by momentary unmasking of the competing audio, in line with ‘glimpsing’ theories of processing speech in noise.

These results open a new window into understanding the dynamics of attention as they wax and wane over time, and the different listening patterns employed for dealing with the influx of sensory input in multisensory environments. Moreover, the novel approach developed here for tracking the locus of momentary attention in a naturalistic virtual-reality environment holds high promise for extending the study of human behavior and cognition and bridging the gap between the laboratory and real-life.

## Introduction

Focusing attention on one speaker in a noisy environment can be challenging, particularly on the background of other irrelevant speech (McDermott, 2009). Despite the difficulty of this task, comprehension of an attended speaker is generally good and the content of distractor speech is rarely recalled explicitly (Cherry, 1953; Lachter et al., 2004). Preferential encoding of attended speech in multi-speaker contexts is also mirrored by enhanced neural responses to attended vs. distractor speech (Ding and Simon, 2012; Mesgarani and Chang, 2012; O’Sullivan et al., 2015; Zion Golumbic et al., 2013b). However, there are also indications that distractor speech is processed, at least to some degree. Examples for this are the Irrelevant Stimulus Effect, where distractor words exert priming effect on an attended task (Beaman et al., 2007; Neely and LeCompte, 1999; Treisman, 1964), as well as occasional explicit detection of salient words in distractor streams (Cherry, 1953; Parmentier et al., 2018; Röer et al., 2017; Wood and Cowan, 1995). These effects highlight a key theoretical tension regarding how processing resources are allocated among competing speech inputs. Whereas Late-Selection models of attention posit that attended and distractor speech can be fully processed, allowing for explicit detection of words in so-called unattended speech (Deutsch and Deutsch, 1963; Duncan, 1980; Parmentier et al., 2018), Limited-Resources models hold that there are inherent bottlenecks for linguistic processing of concurrent speech due to limited resources (Broadbent, 1958; Lachter et al., 2004; Lavie et al., 2004; Raveh and Lavie, 2015). The latter perspective reconciles indications for occasional processing of distractor speech as stemming from rapid shifts of attention toward distractor speech (Conway et al., 2001; Escera et al., 2003; Lachter et al., 2004). Yet, despite the parsimonious appeal of this explanation, to date there is little empirical evidence supporting and characterizing the psychological reality of attention switches among concurrent speakers.

Establishing whether and when rapid shifts of attention towards distractor stimuli occur is operationally challenging since it refers to individuals’ internal state that researchers do not have direct access to. Existing metrics for detecting shifts of attention among concurrent speech primarily rely on indirect measures such as prolongation of reaction times on an attended task (Beaman et al., 2007) or subjective reports (Wood and Cowan, 1995). Given these limitations, current understanding of the dynamics of attention over time, and the nature and consequences of rapid attention-shifts among concurrent speech, is extremely poor. Nonetheless, gaining insight into the dynamics of internal attention-shifts is critical for understanding how attention operates in naturalistic multi-speaker settings.

Here we introduce a new experimental platform for studying the dynamic deployment of attention among concurrent speakers. We utilize Virtual Reality (VR) technology to simulate a naturalistic audio-visual multi-speaker environment, and track participants’ gaze-position within the Virtual Scene as a marker for the locus of overt attention and as a means for detecting attention-shifts among concurrent speakers. Participants experienced sitting in a “Virtual Café” across from a partner (avatar; animated target speaker) and were required to focus attention exclusively towards this speaker. Additional distracting speakers were placed at surrounding tables, with their number and location manipulated across conditions. Continuous tracking of gaze-location allowed us to characterize whether participants stayed focused on the target speaker as instructed or whether and how often they performed overt glimpses around the environment and toward distractor speakers. We further tested whether gaze-shifts are associated with salient acoustic changes in the environment, such as onsets in distractor speech that can potentially grab attention exogenously (Wood and Cowan, 1995) or brief pauses that create momentary unmasking of competing sounds (Cooke, 2006; Lavie et al., 2004).

Gaze-shifts are often used as a proxy for attention shifts in natural vision (Anderson et al., 2015; Schomaker et al., 2017; Walker et al., 2017), however this measure has not been utilized extensively in dynamic contexts (’t Hart et al., 2009; Foulsham et al., 2011). This novel approach enabled us to characterize the nature of momentary attention-shifts in ecological multi-speaker listening conditions, as well as individual differences, gaining insight into the factors contributing to dynamic attention shifting and its consequences on speech comprehension.

## Methods

### Participants

26 adults participated in this study (ages 18–32, median 24; 18 female, 3 left handed), all fluent in Hebrew, with normal hearing and no history of psychiatric or neurological disorders. Signed informed consent was obtained from each participant prior to the experiment, in accordance with guidelines of the Institutional Ethics Committee at Bar-Ilan University. Participants were paid for participation or received class credit.

### Apparatus

Participants were seated comfortably in an acoustic-shielded room and viewed a 3D Virtual Reality scene of a café, through a head-mounted device (Oculus Rift Development Kit 2). The device was custom-fitted with an embedded eye-tracker (SMI, Teltow, Germany; 60Hz monocular sampling rate) for continuous monitoring of participants’ eye-gaze position. Audio was presented through high-quality headphone (Sennheiser HD 280 pro).

### Stimuli

Avatar characters were selected from the Mixamo platform (Adobe Systems, San Jose, CA). Soundtracks for the avatars’ speech were 35-50 seconds long segments of natural Hebrew speech taken from podcasts and short stories (www.icast.co.il), and avatars’ mouth and articulation movements were synced to the audio to create a realistic audio-visual experience of speech (LipSync Pro, rogo digital, England). Scene animation and experiment programming was controlled using an open-source VR engine (Unity Software, unity3d.com). Speech audio was manipulated within Unity using a 3D sound algorithm, so that it was perceived as originating from the spatial location of the speaking avatar. Participants’ head movements were not restricted, and both the graphic display and 3D sound were adapted on-line in accordance with head-position, maintaining a spatially-coherent audio-visual experience.

### Experiment design

In the Virtual Café setting, participants experienced sitting at a café table facing a partner (animated speaking avatar) telling a personal narrative. They were tasked to focus attention exclusively on the speech of their partner (target speaker) and to subsequently answer four multiple-choice comprehension questions about the narrative (e.g. “What computer operating system was mentioned?”). Additional pairs of distracting speakers (avatars) were placed at surrounding tables, and we systematically manipulated the number and location of distractors in four conditions: No Distraction (NoD), Left Distractors (LD), Right Distractors (RD), Right and Left Distractors (RLD; Figure 1). Each condition consisted of five trials (∼4 minutes per condition), and their order was randomized. The identity and voice of the main speaker was kept constant throughout the experiment, with different narratives in each trial, while the avatars and narratives serving as distractors varied from trial to trial. The allocation of each narrative to condition was counter-balanced across participants, to avoid material-specific biases. A training session was performed at the beginning of the experiment to familiarize participants with the environment and the type of comprehension questions asked.

**Figure 1:**
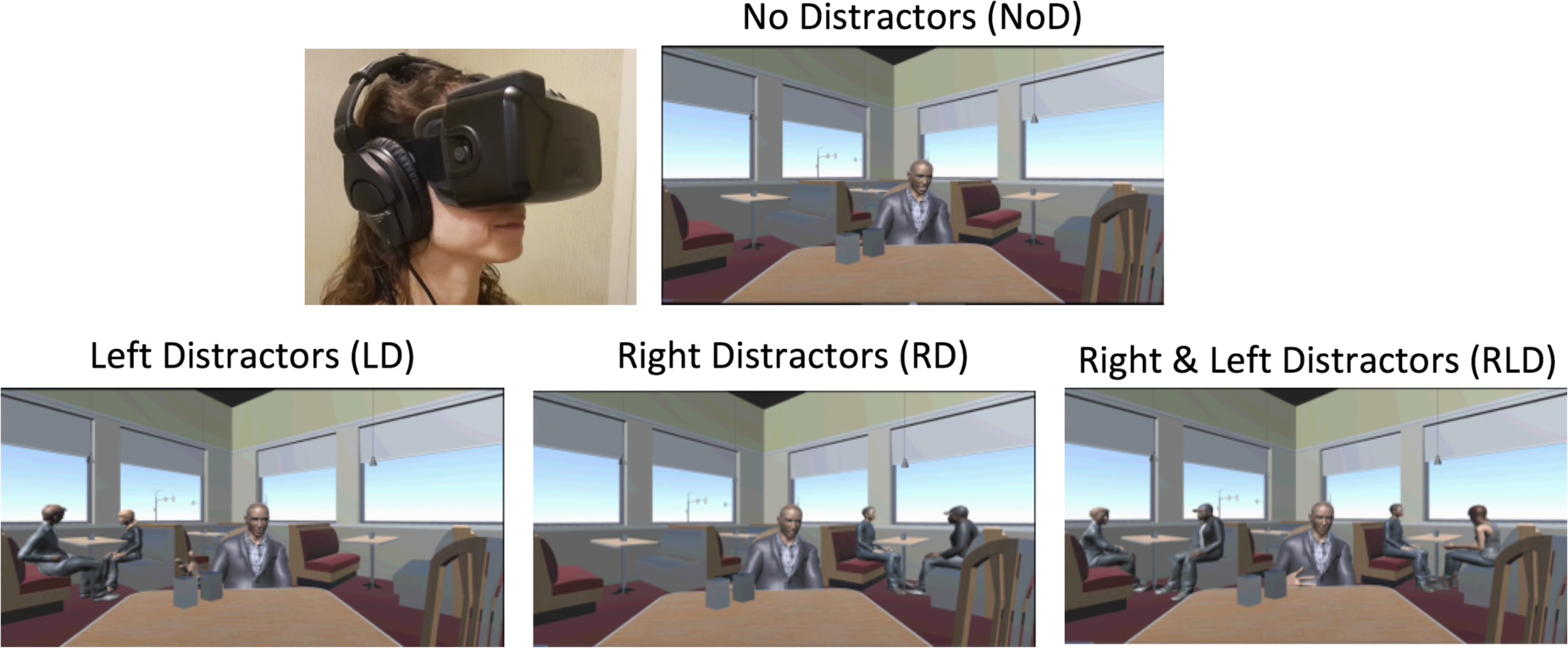
Examples of the four conditions encountered in the Virtual Café. Participants are instructed to attend to the narrative of the target speaker facing them. The number and location of distractor speakers was manipulated across conditions

### Analysis of Eye-Gaze Dynamics

Analysis of eye-gaze data was performed in Matlab, (Mathworks, Natick, MA) using functions from the fieldtrip toolbox (fieldtriptoolbox.org) as well as custom-written scripts. The position of eye-gaze position in virtual space coordinates (x,y,z) was monitored continuously throughout the experiment. Periods surrounding eye-blinks were removed from the data (250ms around each blink). Clean data from each trial was analyzed as follows:

First, we mapped gaze-positions onto specific avatars/locations in the 3D virtual scene. For data reduction, we used a spatial clustering algorithm (k-means) to combine gaze data-points associated with similar locations in space. Next, each spatial cluster was associated with the closest avatar, by calculating the Euclidean distance between the center of the cluster and the center of each avatar presented in that condition. If two or more clusters were associated with looking at the same avatar, they were combined. Similarly, clusters associated with the members of the distractor avatar-pairs (left or right distractors) were combined. If a cluster did not fall within a particular distance-threshold from any of the avatars, it was associated with looking at “the Environment”. This resulted in a maximum of four clusters capturing the different possible gaze locations in each trial: (1) Target Speaker, (2) Left Distractors (when relevant), (3) Right Distractors (when relevant) and (4) Rest of the Environment. The appropriateness of cluster-to-avatar association and distance-threshold selection was verified through visual inspection.

Based on the clustered data, we quantified the percent of time that participants spent focusing at each location (*Percent Gaze Time*) in each trial, and detected the times of *Gaze-Shifts* from one cluster to another. The number of Gaze-shifts as well as the Percent Gaze Time spent at each of the four locations - Target Speaker, Left Distractors, Right Distractors and Environment – were averaged across trials, within condition. Since conditions differed in the type and number of distractors, comparison across conditions focused mainly on metrics pertaining to gazing at / away-from the target speaker. One-way ANOVAs with repeated measures were performed to test whether the Percent Gaze Time towards the target speaker or the number of Gaze-shifts away from the target speaker were modulated by the type of distraction (NoD, LD, RD, RLD). In addition, in order to assess the effect of eye-gaze location on speech comprehension we calculated the Pearson correlation between performance on the comprehension questions and the percent time spent looking at the target speaker, across participants.

#### Analysis of Speech Acoustics relative to Gaze-Shifts

A key question is what prompts overt gaze-shifts away from the target speakers, and specifically whether they are driven by changes in the acoustic input or if they should be considered more internally-driven. Two acoustic factors that have been suggested as inviting attention-shifts among concurrent speech are (a) onsets / loudness increases in distractor speech that can potentially grab attention exogenously (Wood and Cowan, 1995), and (b) brief pauses that create momentary unmasking of competing sounds (Cooke, 2006; Lavie et al., 2004). To test whether one or both of these factors account of occurrence of gaze-shifts away from the target speaker in the current data, we performed a gaze-shift time-locked analysis of the speech-acoustics of target speech (in all conditions) and distractor speech (in the LD, RD and RLD conditions).

To this end, we first calculated the temporal envelope of the speech presented in each trial using a windowed RMS (30 ms smoothing). The envelopes were segmented relative to the times where gaze-shifts *away from the target speaker* occurred in that particular trial (− 400 to + 200 ms around each shift). Since the number of gaze-shifts varied substantially across participants, we averaged the gaze-shift-locked envelope-segments across all trials and participants, within condition. The resulting average acoustic-loudness waveform in each condition was compared to a distribution of non-gaze-locked loudness levels, generated through a permutation procedure as follows: The same acoustic envelopes were segmented randomly into an equal number of segments as the number of gaze-shifts in each condition (sampled across participants with the same proportion as the real data). These were averaged, producing a non-gaze-locked average waveform. This procedure was repeated 1,000 times and the real gaze-shift locked waveform was compared to the distribution of non-gaze-locked waveforms. We identified time-points where the loudness level fell above or below the top/bottom 5%^tile^ of the non-gaze-locked distribution, signifying that the speech acoustics were particularly quiet or loud relative (relative to the rest of the presented speech stimuli).

## Results

### Behavioral results

Accuracy on the multiple-choice comprehension questions of the target speaker was relatively good (mean accuracy 82%±3 across all conditions). Accuracy rates did not differ significantly across conditions [F(3,75) =2.206, p=0.119; Figure 2].

**Figure 2:**
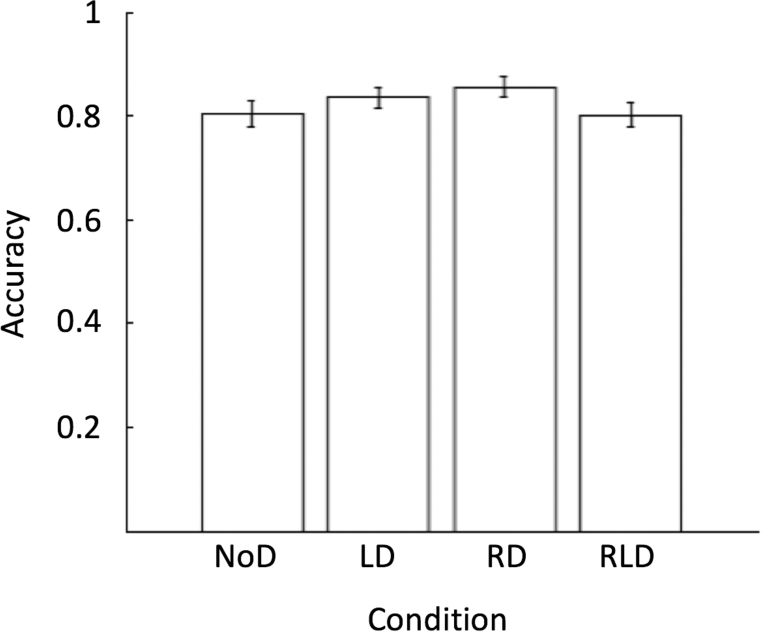
Accuracy rates on comprehension questions across conditions. Error bars indicate Standard Error of the Mean (SEM).

### Eye-Gaze Patterns

#### No effect of audio-visual distraction on gaze patterns

Figure 3 shows the average *Percent Gaze Time* away from the target speaker (i.e., time spent looking toward distractor avatars or other locations in the Environment) as well as the number of *gaze-shifts* away from the target speaker, across conditions. On average, participants spent ∼7.6% of each trial (∼3 seconds in a 40-second-long trial) looking at locations other than the target speaker and they performed an average of 2.5 gaze-shifts per trial. The average number of gaze-shifts and percent gaze-time away from the target speaker did not differ significantly across conditions [percent gaze-time away: F(3,75) = 1.286, p= 0.285; number of gaze-shifts: F(3,75) = 1.722, p= 0.185], suggesting that these were not affected by the presence or severity of audio-visual distraction.

**Figure 3:**
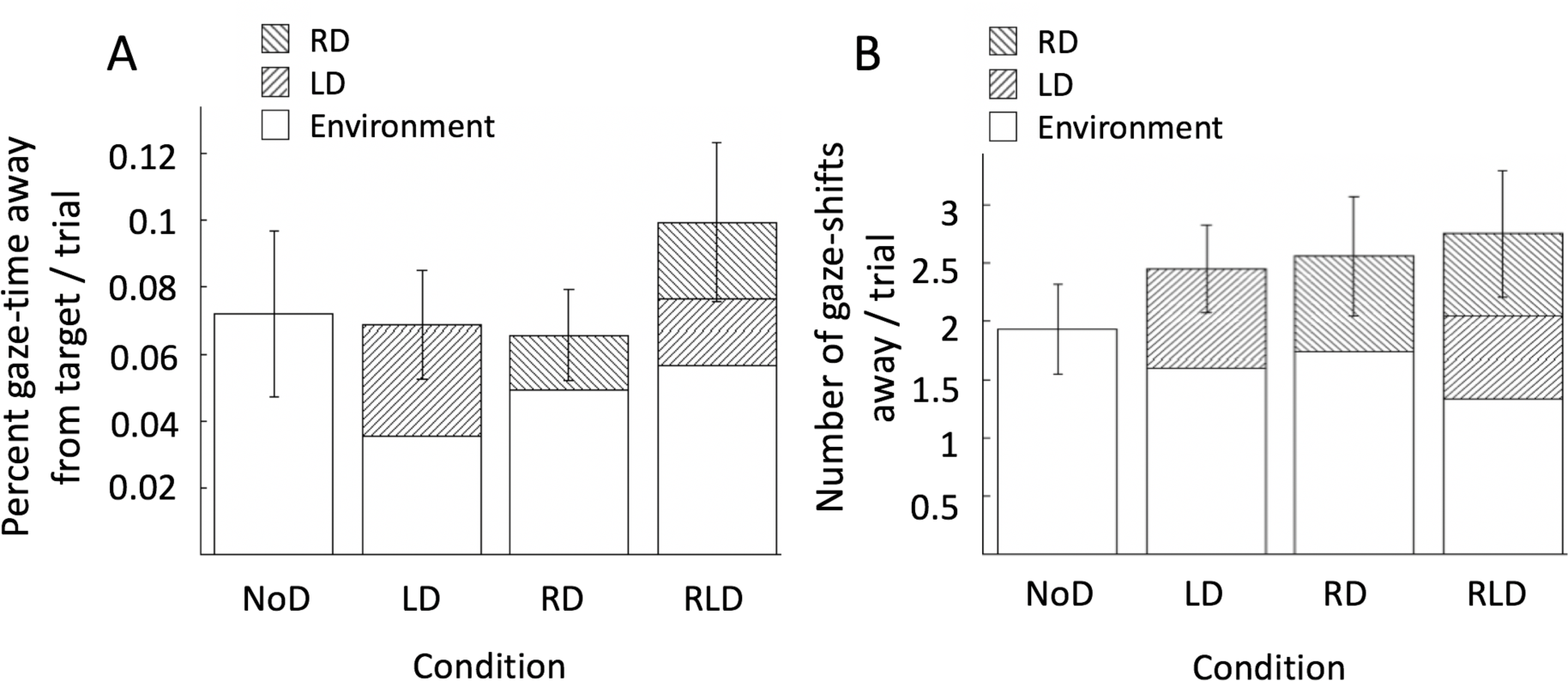
Summary of gaze-shift results away from the target speaker across conditions. (A) Proportion of Gaze-Time and (B) Number of gaze-shifts per trial away from target speaker. Results within each condition are broken down by gaze-location (Right Distractors, Left Distractors or Environment in blank, left and right diagonals, respectively). Error bars indicate Standard Error of the Mean (SEM). There was no significant difference between conditions in the total Gaze-time away from the target speaker or number of gaze-shifts.

### Individual Differences in Gaze Patterns and Link to Behavior

Despite the lack of differences in gaze-shift frequency across distraction conditions at the group-level, there was substantial variability between participants in the number of gaze-shift performed and percent time spent gazing away from the target speaker. As illustrated in Figures 4 & 5, some participants stayed completely focused on the main speaker throughout the entire experiment, whereas others spent a substantial portion of each trial gazing around the environment (*range across participants:* 0-18 average number gaze-shifts per trial; 0-34.52% average percent of trial spent looking away from the target speaker). This motivated further inspection of gaze-shift behavior at the individual level. Specifically, we tested whether individual behavior of performing many or few gaze-shifts away from the target were stable across conditions. We calculated Cronbach’s Alpha between conditions and found high internal consistency across conditions in the number of gaze-shifts performed as well as in the percent of gaze-time away from the target speaker (alpha = 0.889 and alpha = 0.832 respectively). This was further demonstrated by a significant positive correlation between the mean time spend gazing away from the target speaker in the two extreme conditions: No Distraction vs. Right & Left Distraction (r = 0.7879, p<10^− 5^ and r=0.7544, p< 10^− 5^ respectively; Figure 5C&D). This pattern suggests that individuals have characteristic tendencies to either stay focused or gaze-around the scene, above and beyond the specific sensory attributes or degree of distraction in a particular scenario.

**Figure 4:**
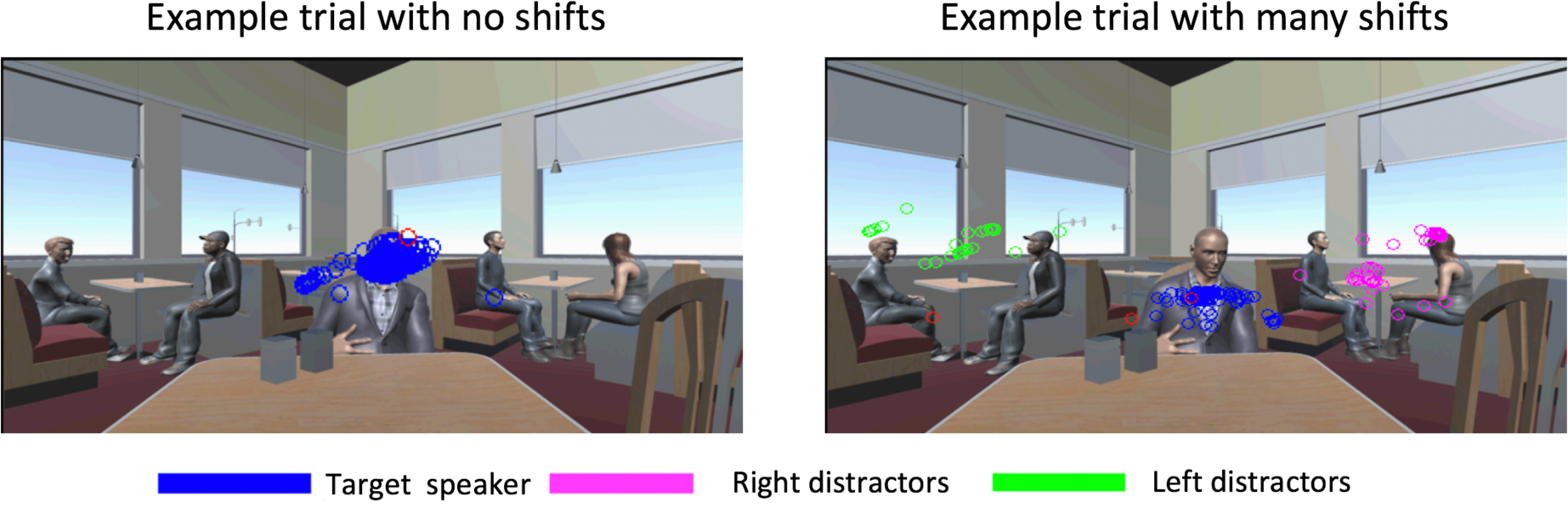
Illustration of the variability in gaze-patterns across individuals. The figure depicts all gaze data points in a specific trial in the RLD condition for two example participants. While the participant shown in the left panel remained focused exclusively on the target speaker throughout the trial (blue dots), the participant in the right panel spent a substantial portion of the trial looking at the distractor speakers on both the left (green) and the right (magenta).

**Figure 5:**
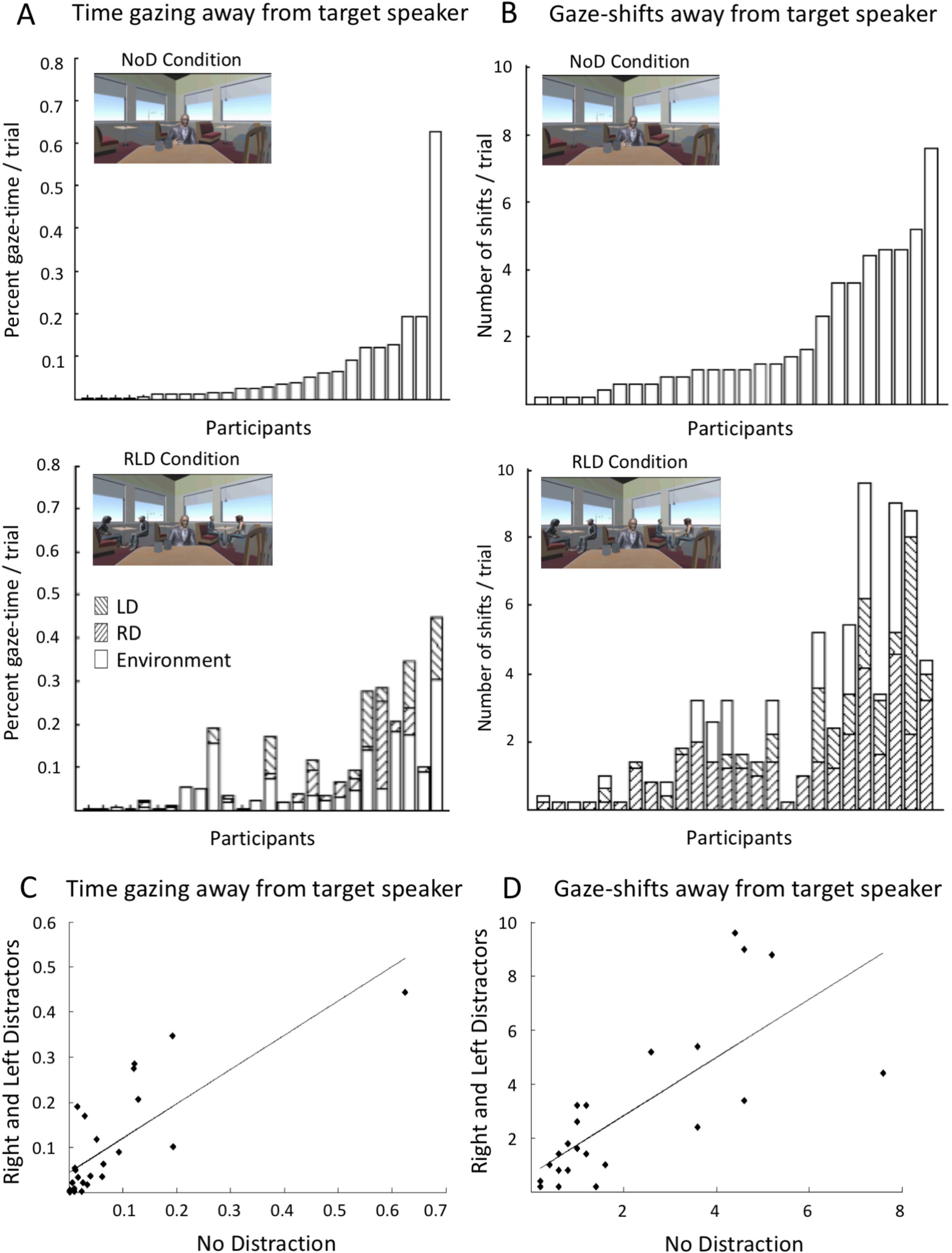
Individual gaze-shift patterns. **(A):** Percent of time spent gazing away from the target speaker and (**B)** average umber of gaze-shifts per trial in the NoD condition (top) and the RLD conditions (bottom), across individual participants. In both cases, participant order is sorted by the NoD condition (top panels). **(C)** Pearson’s correlation of the percent of time pent gazing away from the target speaker and (**D)** average number of gaze-shifts per trial, between the two extreme onditions: NoD vs. RLD

Next, we tested whether shifting ones’ gaze away from the target speaker impacted speech comprehension. The number of gaze-shift was not significantly correlated with accuracy rates on the comprehension questions (p=0.39), however the mean time spent gazing away from the target speaker was significantly correlated with comprehension performance (r = −0.5496, p < 0.004; Figure 6), suggesting that spending more time looking around the environment had negative consequences on comprehension of the target speaker.

**Figure 6:**
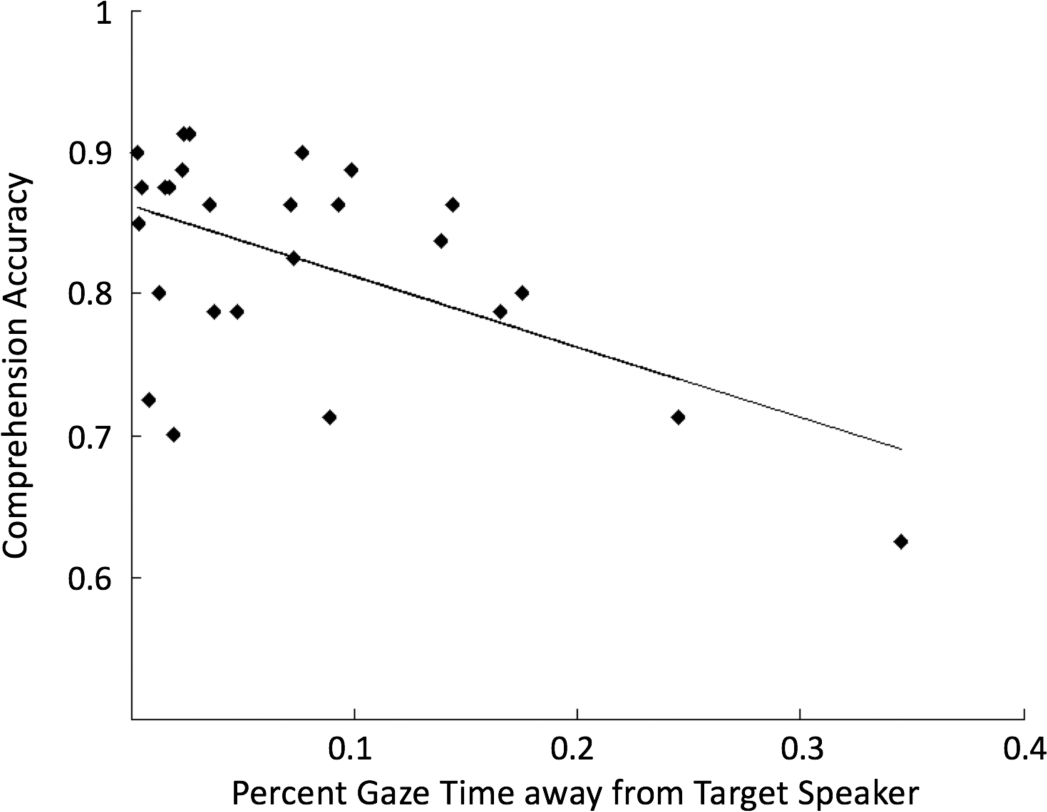
Correlation between the time spent gazing away from target speaker, averaged across all conditions, vs. performance accuracy on the comprehension task

#### Gaze-locked analysis of speech acoustics

Last, we tested whether there was any relationship between the timing of gaze-shifts and the local speech-acoustics. To this end, we performed a gaze-shift-locked analysis of the envelope of the target or distractor speech (when present). Analysis of distractor speech envelope consisted only of eye-gaze shifts *toward that distractor* (i.e. excluding shifts to other places in the environment). Figure 7 shows the average time-course of the target and distractor speech envelopes relative to the onset of a gaze-shift. For both target speech (top row) as well as for distractor speech (bottom row), gaze-shifts seem to have been preceded by a brief period of silence (within the lower 5%^tile^; red shading) between 200-300ms prior to the shift. This is in line with an acoustic release-from-masking account, suggesting that gaze-shifts are prompted by momentary gaps in the speech, and particularly when gaps in concurrent speech coincide-temporally (as seen here in the Single and Two Distractor conditions). Conversely, the suggestion that attention-shifts are a product of exogenous capture by salient events in distracting speech does not seem to be supported by the current data, since the acoustics of the distractor speech that participants shifted their gaze towards did not seem to contain periods with consistently loud acoustics. We did however find increases in loudness of the target speech acoustics near gaze-shift onset (within the top 5%^tile^; red shading between −100 to +50ms).

**Figure 7:**
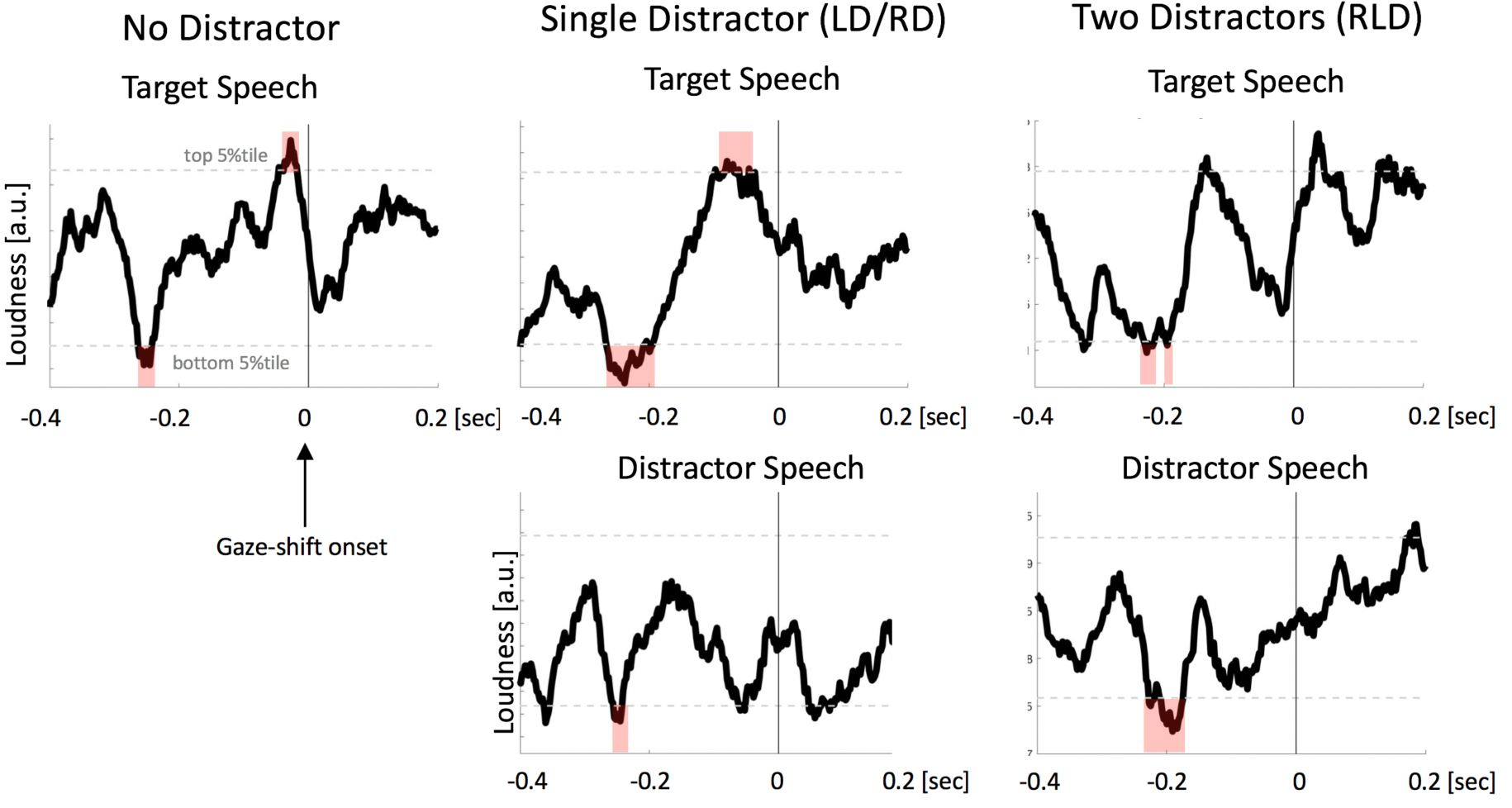
Gaze-shift locked analysis of speech acoustics. Average time-course of the target (top) and distractor bottom) speech envelopes relative to gaze shift onset (t=0). Horizontal dotted gray lines depict the top and bottom %tile of loudness values generated through the permutation procedure of non-gaze-locked acoustics segments. The shaded red areas indicate time-periods where the speech sound-level fell within the lower/upper 5%^tile^ of the istribution, respectively.

## Discussion

The current study is a first and novel attempt to characterize how individuals deploy overt attention in naturalistic audiovisual settings, laden with rich and competing stimuli. By monitoring eye-gaze dynamics in our Virtual Café, we studied the patterns of gaze-shifts and its consequences for speech comprehension. Interestingly, we found that the presence and number of competing speakers in the environment did <u>not</u>, on average, affect the frequency of gaze-shifts away from the target speaker, nor did it impair comprehension of the target speaker. Rather, our results suggest that gaze-control under these conditions is highly individualized. Participants displayed characteristic patterns of either staying focused on a target speaker or sampling other locations in the environment overtly, regardless of the severity of the so-called sensory distraction. Critically, the amount of time that individuals spent looking around the environment and away from the target speaker was negatively correlated with speech comprehension, directly linking overt attention to speech comprehension. We also found that gaze-shifts away from the target speaker occurred primarily during gaps in the acoustic input, suggesting they are prompted by momentary unmasking of the competing audio, in line with ‘glimpsing’ theories of processing speech in noise. These results open a new window into understanding the dynamics of attention as they wax and wane over time, and the listening patterns exhibited by individuals for dealing with the influx of sensory input in complex naturalistic environments.

### Is Attention Stationary?

An underlying assumption of many experimental studies is that participants allocate attention solely to task-relevant stimuli, and that attention remains stationary over time. However this assumption is probably unwarranted (Esterman et al., 2013; Weissman et al., 2006) since sustaining attention over long periods of time is extremely taxing (Avisar and Shalev, 2011; Schweizer and Moosbrugger, 2004; Warm et al., 2008), and individuals spend a large proportion of the time mind-wandering or ‘off-task’ (Boudewyn & Carter, 2018; Killingsworth & Gilbert, 2010; but cf. Seli et al., 2018). Yet, empirically testing the studying the frequency and characteristics of attention shifts is operationally difficult since it pertains to participants’ internal state that experimenters don’t have direct access to. The use of eye-gaze position as a continuous metric for the locus of momentary overt attention in a dynamic scene in the current study contributes to this endeavor.

We find that roughly 30% of participants spent over 10% of each trial looking at places in the environment other than the to-be-attended speaker, across all conditions. Interestingly, this proportion is similar to that reported in previous studies for the prevalence of detecting ones’ own name in a so-called unattended message (Cherry, 1953; Wood & Cowan, 1995), an effect attributed by some to rapid attention shifts (Beaman et al., 2007; Lachter et al., 2004; Lin and Yeh, 2014). Although in the current study we did not test whether these participants also gleaned more information from distractors’ speech, we did find that comprehension of the target speaker was reduced in these participants. These results highlight the importance of studying individual differences in attentional control and suggest that this may hold a key for understanding some of the intriguing behavioral effects for partial processing of unattended speech.

In the current study set we did not collect additional personal data from participants which may have shed light on the source of the observed variability in gaze-patterns across individuals. However, based on extensive literature on individual differences in attentional performance, individual differences may stem from factors such as susceptibility to distraction (Avisar and Shalev, 2011; Bourel-Ponchel et al., 2011; Cowan et al., 2005; Ellermeier and Zimmer, 1997; Forster and Lavie, 2014; Hughes, 2014), working memory capacity (Conway et al., 2001; Hughes, 2014; Kane and Engle, 2002; Naveh-Benjamin et al., 2014; Sörqvist et al., 2013; Tsuchida et al., 2012; Wiemers and Redick, 2018) or personality traits (Baranes et al., 2015; Hoppe et al., 2018; Rauthmann et al., 2012; Risko et al., 2012). Additional dedicated research is needed to resolve the source of the individual differences observed here.

### Is eye-gaze a good measure for attention-shifts among concurrent speech?

One may ask, to what extent do the current results fully capture the prevalence of attention-shifts, since it is known that these can also occur covertly (Petersen and Posner, 2012; Posner, 1980)? This is a valid concern and indeed the current results should be taken as representing a *lower-bound* for the frequency of attention-shifts and we should assume that attention-shifts are probably more prevalent than observed here. This motivates the future development of complementary methods for quantifying covert shifts of attention among concurrent speech, given the current absence of a reliable metrics.

Another concern that may be raised with regard to the current results is that individuals may maintain attention to the target speaker even while looking elsewhere, and hence the gaze-shifts measured here might not reflect true shifts of attention. Although in principle this could be possible, previous research shows that this is probably not the default mode of listening under natural audiovisual conditions. Rather, a wealth of studies demonstrate a tight link between gaze-shifts and attention-shifts (Chelazzi et al., 1995; Deubel and Schneider, 1996; Grosbras et al., 2005; Szinte et al., 2018) and gaze is widely utilized experimentally as a proxy for the locus of visuospatial attention (Gredebäck et al., 2009; Linse et al., 2017). In multi-speaker contexts it has been shown that participants tend to move their eyes towards the location of attended speech sounds (Gopher, 1973; Gopher and Kahneman, 1971). Similarly, looking towards the location of distractor-speech significantly reduces intelligibility and memory for attended speech and increases intrusions from distractor speech (Reisberg et al., 1981; Spence et al., 2000; Yi et al., 2013). This is in line with the current finding of a negative correlation between the time spent looking at the target speaker and speech comprehension, further linking overt gaze to selective attention to speech. Studies on audiovisual speech processing further indicate that looking at the talking face increases speech intelligibility and neural selectivity for attended speech (Crosse et al., 2016; Lou et al., 2014; Park et al., 2016; Sumby and Pollack, 1954; Zion Golumbic et al., 2013a), even when the video is not informative about the content of speech (Kim and Davis, 2003; Schwartz et al., 2004), and eye-gaze is particularly utilized for focusing attention to speech under adverse listening condition (Yi et al., 2013). Taken together, this findings support the interpretation that gaze-shifts reflect shifts in attention away from the target speaker, in line with the limited resources perspective of attention (Esterman et al., 2014; Lavie et al., 2004), making eye-gaze a useful and reliable metric for studying the dynamics of attention to naturalistic audio-visual speech. Interestingly, this metric has recently been capitalized on for use in assistive listening devices, utilizing eye-gaze direction to indicate the direction of a listeners attention (Favre-Felix et al., 2017; Kidd, 2017).

### Listening between the Gaps – what prompts attention shifts among concurrent speech?

Besides characterizing the prevalence and behavioral consequences of attention-shifts in audio-visual multi-talker contexts, it is also critical to understand what prompts these shifts. Here we tested whether there are aspects of the scene acoustics that can be associated with attention-shifts away from the target speaker. We specifically tested two hypotheses: (1) that attention is captured exogenously by highly salient sensory events in distracting speech (Itti and Koch, 2000; Kayser et al., 2005; Wood and Cowan, 1995), and (2) that attention-shifts occur during brief pauses in speech acoustics that momentarily unmask the competing sounds (Cooke, 2006; Lavie et al., 2004).

Regarding the first hypothesis, the current data suggest that distractor saliency is not a primary factor in prompting gaze-shifts. Since gaze-shifts were just as prevalent in the NoD condition as in conditions that contained distractors and since no consistent increase in distractor loudness was observed near gaze-shifts, we conclude that the gaze-shifts performed by participants do not necessarily reflect exogenous attentional capture by distractor saliency. This is in line with previous studies suggesting that sensory saliency is less effective in drawing exogenous attention in dynamic scenarios relative to the stationary contexts typically used in laboratory experiments (Smith et al., 2013).

Rather, our current results seem to support the latter hypothesis that attention-shift are prompted by momentary acoustic release-from-masking. We find that gaze-shifts occurred more consistently ∼200-250 ms after instances of <u>low</u> acoustic intensity in both target and distractor sounds, which is on-par with the initiation time for saccades (Gilchrist, 2011). This pattern fits with accounts for comprehension of speech-in-noise, suggesting that listeners utilize brief periods of unmasking or low SNR to glean and piece together information for deciphering speech content (‘acoustic glimpsing’; Cooke, 2006; Li & Loizou, 2007; Rosen, Souza, Ekelund, & Majeed, 2013; Vestergaard, Fyson, & Patterson, 2011). Although this acoustic-glimpsing framework is often used to describe how listeners maintain intelligibility of target-speech in noise, it has not been extensively applied to studying *shifts* of attention among concurrent speech. The current results suggest that brief gaps in the audio or periods of low SNR may serve as triggers for momentary attention shifts, which can manifest overtly (as demonstrate here), and perhaps also covertly. Interestingly, a previous study found that eye-blinks also tend to occur more often around pauses when viewing and listening to audio-visual speech (Nakano and Kitazawa, 2010), pointing to a possible link between acoustic glimpsing and a reset in the oculomotor system, creating optimal conditions for momentary attention-shifts.

## Conclusions

There is growing understanding that in order to really understand the human cognitive system, it needs to be studied in contexts relevant for real-life behavior, and that tightly constrained artificial laboratory paradigms do not always generalize to real-life (’t Hart et al., 2009; Foulsham et al., 2011; Hoppe et al., 2018; Kingstone et al., 2008; Risko et al., 2016; Rochais et al., 2017). The current study represents the attempt to bridge this gap between the laboratory and real-life, by studying how individuals spontaneously deploy overt attention in a naturalist virtual-reality environment. Using this approach, the current study highlights the characteristics and individual differences in selective attention to speech under naturalistic listening conditions. This pioneering work opens up new horizons for studying of how attention operates in real-life and understanding the factors contributing to success as well as the difficulties in paying attention to speech in noisy environments.

